# Measurements of neurite extension and nucleokinesis in an iPSC-derived model system following microtubule perturbation

**DOI:** 10.1101/2024.02.06.579144

**Authors:** Muriel Sébastien, Alexandra L. Paquette, Lilian Ferotin, Adam G. Hendricks, Gary J. Brouhard

## Abstract

In neurons, patterns of different microtubule types are essential for neurite extension and nucleokinesis. Cellular model systems such as rodent primary cultures and induced pluripotent stem cells (iPSC)-derived neurons have provided key insights into how these patterns are created and maintained through the action of microtubule-associated proteins (MAPs), motor proteins, and regulatory enzymes. iPSC-derived models show tremendous promise but lack benchmarking and validation relative to rodent primary cultures. Here we have characterized a recent iPSC-derived model, in which doxycycline-induced expression of Neurogenin-2 drives consistent trans-differentiation into the neuronal state (EBiSC-NEUR1 neurons, referred to as NGN2 neurons below). We developed a suite of open-access, semi-automated methods to measure neurite extension and nucleokinesis of NGN2 neurons, which compare favorably to published data from other models. Then, we challenged NGN2 neurons with a panel of drugs that perturb microtubule physiology. NGN2 neurons extension and nucleokinesis were significantly perturbed by two microtubule-targeting drugs, namely a taxane (paclitaxel) and a vinca alkaloid (DZ-2384). In contrast, inhibition of microtubule severing (spastazoline) or of deacetylation (trichostatin A) had a limited effect on nucleokinesis only. Our results support the primary importance of microtubule dynamics in neuronal development and demonstrate the power of NGN2 neurons as a model system.

## Introduction

More than 60 years after the discovery of microtubules in plant cells (*1*), researchers continue to discover new and diverse characteristics of these indispensable polymers. Key among these characteristics is dynamic instability (*2*), or the capacity of microtubule ends to switch rapidly between growth and shrinkage (“catastrophes” and “rescues”). Cells can increase the rates of this switching behavior, e.g., when microtubule plus ends become highly dynamic during mitosis (*3-6*). Alternatively, cells can suppress dynamics, creating microtubules that are long-lived, stable, and resistant to mechanical damage (*4, 7-9*). Both stable and dynamic subsets of microtubules exist side-by-side and are found in diverse geometric arrangements (*10*). These complex networks are built and maintained by a host of micro-tubule-associated proteins (MAPs) (*11*), motor proteins (*12, 13*), and regulatory enzymes (*14*). The coordinated action of these MAPs, motors, and enzymes will construct and functionalize a microtubule cytoskeleton that meets each differentiated cell’s needs (*15*).

Neurons have evolved a particularly sophisticated microtubule network, given their need to extend many neurites, functionalize an axon and dendrites, and transport cargos over great distances (*16, 17*). One of such cargo is the nucleus which will be translocated during neuronal development, a process called nucleokinesis. As neurons develop, they express several isoforms of α- and β-tubulin, including a neuronal specific β-tubulin (β3-tubulin, Tubb3 in humans) (*18*). Many of these isoforms are implicated in brain disease when mutated (*19, 20*). In addition, neurons accumulate several post-translational modifications (PTMs; one component of the “tubulin code” (*21)*), such as polyglutamate chains on αβ-tubulin’s disordered C-termini, and acetylation on α-tubulin in the microtubule lumen. Remarkably, distinct PTMs can be found in anti-parallel bundles, an arrangement that may play a critical role in cargo sorting (*22*). Patterns of neuronal microtubules can also be distinguished by their points of origin, which include the centrosome early in development (*23*) as well as acentrosomal sites such as Golgi outposts (*24, 25*). The centrosomal microtubules, which participate in nucleokinesis, are necessary for radial migration of cortical neurons, but not for axon formation (*26, 27*). These observations indicate distinct roles for centrosomal vs. acentrosomal microtubules in the life cycle of these neurons. Presumably then, microtubules within neurons are distinct at the molecular level, in terms of their relative dynamics, their specific cohort of MAPs, their range of PTMs, etc. But we cannot yet draw a complete molecular map of how microtubule subsets emerge, maintain distinction, and subsequently drive neuronal development (*17*).

To study neurite extension and nucleokinesis at the molecular level, model systems include primary cultures from rodents (*28, 29*), model organisms such as C. elegans and D. melanogaster (*30, 31*), and recently developed models based on human induced pluripotent stem cells (iPSCs) (*32*). The use of human iPSCs shows promise for disease modeling (*33, 34*), particularly when combined with CRISPR/Cas9 genome editing techniques (*35-37*). For example, CRISPR-modified iPSCs could be induced first into neural progenitor cells (NPCs) using small molecules like SMAD inhibitors (later referred to as SMADi (38)) and then differentiated into neuronal subtypes with specific growth factor supplementation (*39*). We recently developed a model for X-linked lissencephaly using SMADi-inducible iPSCs that carry mutations in the DCX gene and were differentiated into cortical neurons (*40*). The induction into NPCs with SMADi requires approximately 30 days of time and media, however. More recently, trans-differentiation approaches were developed wherein iPSCs were engineered to express the neural lineage transcription factor Neurog-enin-2 (NGN2) under a doxycycline inducible promotor (*41-44*). Doxycycline treatment drives consistent induction of all iPSCs into neural lineage in only two days. Then, terminal differentiation into specific neuronal subtypes can be achieved as before, using specific factors. Multiple studies investigating microtubule dynamics and cargo transport using these neurons demonstrated their promise for studying the microtubule network (*45-47*), but their physiology remains incompletely characterized.

In the present study, we developed automated measurement and tracking methods to characterize neurite extension and nucleokinesis in iPSCs-derived neurons, namely a fast-inducible NGN2 iPSC line (*41*) obtained from the European Bank of induced pluripotent Stem Cells (EBiSC-NEUR1, hereafter referred to as NGN2 neurons). These methods enabled the generation of a large data set (n=25508 migrating neurons, *n*=14579 fixed neurons), which we present as a resource for analysis and bench-marking. We find that NGN2 neurons extend neurites at rates comparable to rodent primary cultures and other iPSC models, and NGN2 nuclei show robust translocation and migratory behavior. We challenged NGN2 neurons with a panel of small molecules that disrupt microtubule dynamics, target a microtubule severing enzyme, or alter the balance of tubulin acetylation. We find that microtubule patterning is indeed essential for neurite extension and nucleokinesis in NGN2 neurons (as in other models), but more importantly that our automated quantification of these phenomenon is valid and can reveal drug-specific phenotypes efficiently. Overall, our work contributes to the development of iPSC-based cellular models for the neuronal microtubule cytoskeleton.

## Results and discussion

### NGN2 neurons extend an arbor of neurites enriched in β3-tubulin

We began with a baseline characterization of NGN2 neurons differentiated into cortical neurons. These neurons were obtained from induced Neurogenin2 expression with doxycycline for 2 days prior to long-term cryopreservation; aliquots of frozen cells were thawed directly into terminal differentiation media supplemented with doxycycline at t=0 of the experiment (Fig. 1A). To characterize the development of NGN2 neurons over the next 7 days, we examined their morphology and microtubule content using a suite of automated methods, which we have made open access. In nearly every cell, we observed robust development of neuronal processes, an elongated morphology, and positive staining for the neuronal-specific β3-tubulin, Tubb3, as expected (*48, 49*). In contrast, SMADi-inducible cultures sometimes contain non-neuronal cells and/or proliferating NPCs, even in terminal differentiation media, and thus require additional quality control and care (*40*). Figure 1B shows an immunocytochemistry image of Tubb3 staining at day 3 and day 7, and the length of neurites increased significantly over this time period. To quantify this increase, we skeletonized the tubulin images and measured the total length of the neurite network per cell (or arborization length, see Methods). The arborization length increased from 203.7 ± 5.2 µm at 3 days to 433.2 ± 16.7 µm at 7 days (Fig. 1B, mean ± SEM, p<0.0001, ***). This increase in arborization lengths is comparable to lengths measured in rodent neurons (*50-52*) and in other iPSC models (*40, 53*), which is on the order of a hundred micrometers per day (Cf. Table 1). Increased arborization is typically accompanied by increased expression of Tubb3 and increased PTM levels. To confirm these predictions, we performed Western blots using antibodies against Tubb3 and acetylated tubulin (Acetyl). Both antibody signals increased significantly from day 3 to day 7, with an approximately 30% increase in Tubb3 signal and a 70% increase in Acetyl signal over that time period (Fig 1C).

**Table 1:** Arborization lengths in various models. Arborization length (µm/cell) in different study models with references and date of publication. “∼” was used when values were extrapolated from multiple conditions or visually assessed from a graphical representation of the data. Arborization lengths were measured at DIV3 unless specified otherwise.

| Arborization length ($\mu\text{m}/\text{cell}$) | Model | References |
| --- | --- | --- |
| 165 | Rat hypothalamic neurons DIV4 | 1991 (50) |
| ~1000 | Mouse sensory neurons | 1995 (51) |
| 213 | Rat hippocampal neurons | 2004 (52) |
| ~700 | Rat cortical neurons | 2005 (54) |
| ~300 | Human iPSC-derived cortical neurons | 2017 (53) |
| 155 | Human iPSC-derived cortical neurons | 2021 (55) |
| 154 | Human iPSC-derived cortical neurons | 2024 (40) |
| 203 | Human NGN2-induced neurons | This study |

**Figure 1:**
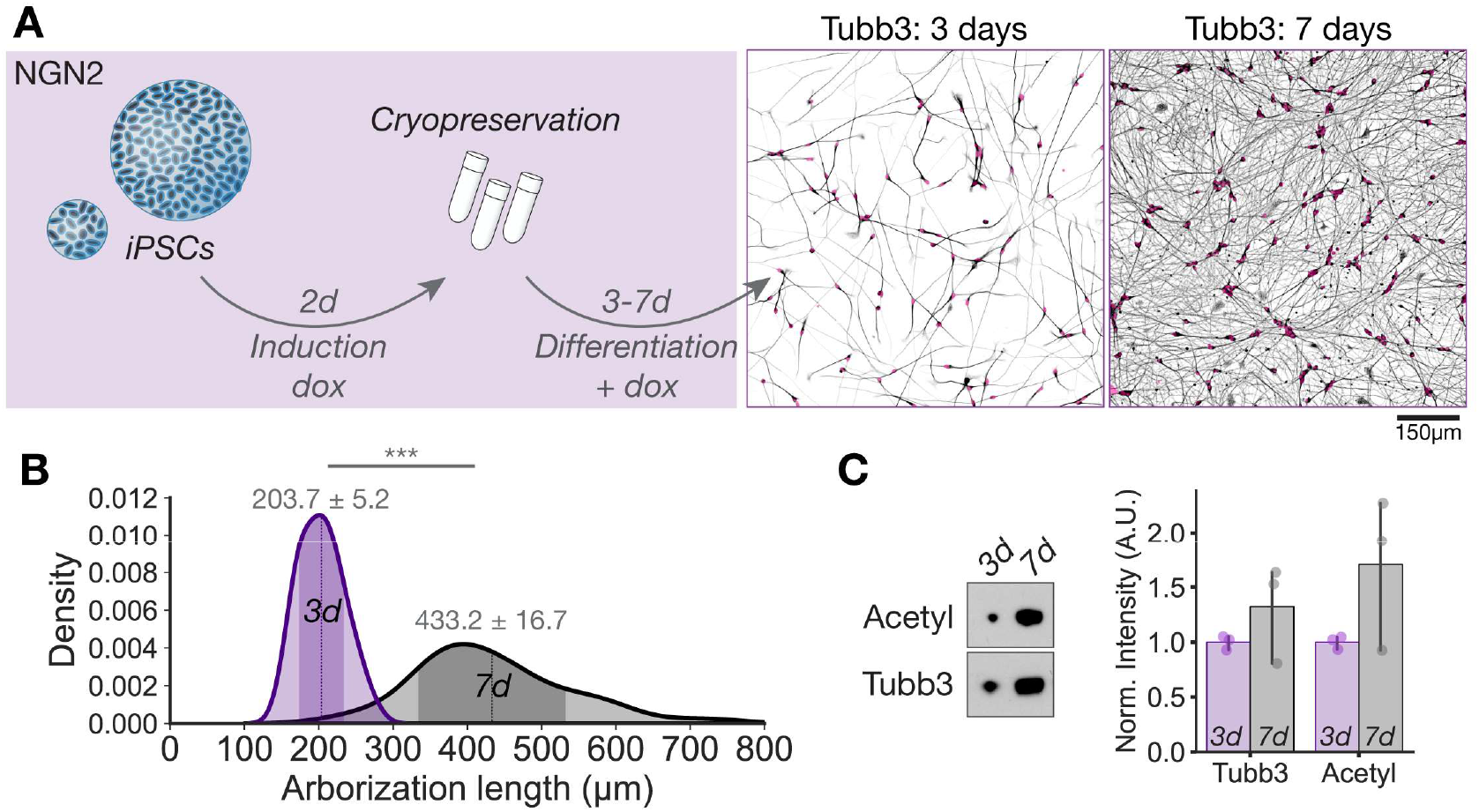
NGN2 neurons extend an arbor of neurites enriched in β3-tubulin. (A) Left panel: Schematics of induction and differentiation process in NGN2 lines from iPSC aggregates to early neuronal stages. NGN2 iPSCs were induced for 2 days in Neurobasal media supplemented with doxycycline (dox) before cryopreservation, shipping and usage. The early neurons were thawed in BrainPhys + dox and terminally differentiated for 3-7 days. Right panels: Representative images of Tubb3 immunostaining of NGN2 neurons at 3 days and 7 days of differentiation. Nuclei are pseudo-colored in magenta. (B) Plot of the distribution of neurite arborization length per cell body in NGN2 neurons at 3 days (3d) and 7 days (7d) of differentiation. *n*=868-4181 neurons from at least 2 independent experiments. The mean values (dotted lines) are detailed on the plot for each condition ± SEM, while the darker areas show standard deviations. Statistical analysis was done using Mann-Whitney non-parametric tests: *** p<0.001. (C) Representative immunoblots and quantifications of tubulin acetylation (611B1, Acetyl) and β3-Tubulin (Tubb3) in NGN2 neurons at 3 (3d) and 7 days (7d) of terminal differentiation.

From these findings, we conclude that NGN2 iPSCs differentiate and mature into cortical neurons that compare favorably to rodent cultures and other iPSC models. In particular, the relative homogeneity of NGN2 neurons make them well-suited to screening applications and automated image analysis, e.g. of cell morphology and nucleokinesis.

### NGN2 neurons exhibit robust nucleokinesis

Nucleokinesis is also a critical process in neuronal development, so we analyzed NGN2 neurons using a previously established nuclear translocation assay (*40*). The cells were seeded at a density of 15000 cells/cm2, differentiated for 3 days, stained with Syto™ nuclear dye, and imaged for 5h with 5min intervals between frames (see Supplemental Movies). We observed robust nuclear movements over the full time-course of imaging. Figure 2A and 2B show representative still images of the cells’ morphologies and nuclear staining, with nuclear trajectories overlayed as purple lines. We quantified nuclear movements using an automated tracking pipeline (*40*). The nuclei were detected and linked throughout frames within TrackMate (*56, 57*) using the Laplacian of Gaussian (LoG) and Linear Assignment Problem (LAP) algorithms respectively (see Methods). Trajectories were filtered and analyzed to determine the mean speed, the total distance traveled, and directionality index. Figure 2C shows a sample of these extracted trajectories (n=1764), which are color-coded according to their total displacement (see scale on the right of Fig. 2C). NGN2 nuclei covered considerable distance over the 5 h imaging period (Fig. 2E) and moved slightly faster (0.43 ± 0.01 µm/min, Fig. 2D) than migrating iPSC-derived neurons (*40, 58*). The directionality index was 0.26 ± 0.01, however, implying that movements are more meandering than directed (Fig. 2F). In addition, we measured the aspect ratio of the nuclei in DAPI stainings, and found that NGN2 nuclei were slightly elongated (Fig. 2G), possibly due to cytoskeletal forces exerted on the nucleus during nucleokinesis.

**Figure 2:**
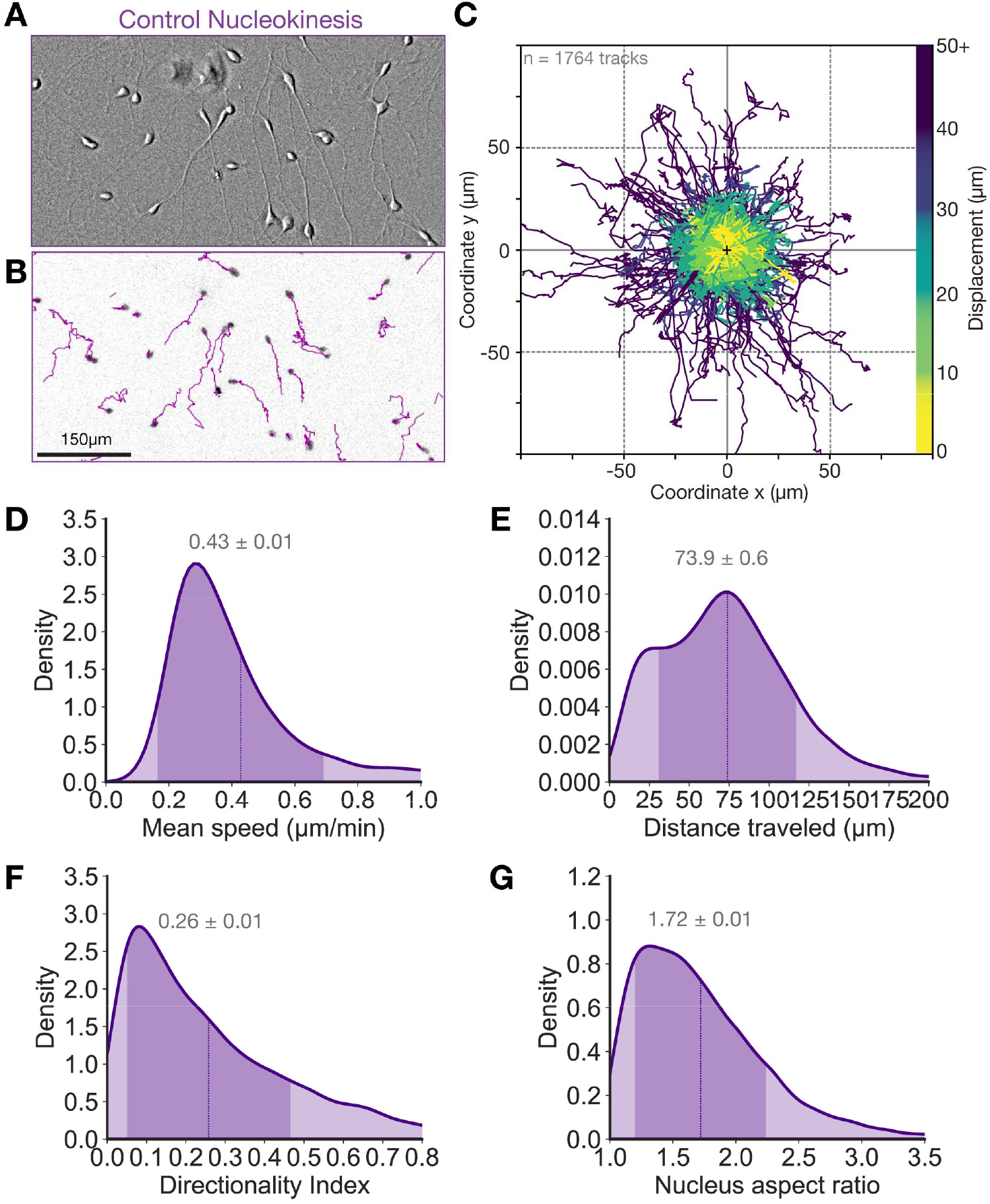
NGN2 neurons exhibit robust nucleokinesis. (A) Representative image of NGN2 neurons at 3 days of differentiation with brightfield imaging, showing neuronal morphology at the start of the tracking experiment. (B) Still image of nuclei stained with 250nM Syto™ deep red for the neurons shown in panel A. Results of the automated tracking over a 5h-time period are showed as overlayed tracks in purple. (C) Position plots of 1764 sample tracks for NGN2 nuclei. Each track is color-coded according to its displacement (see color scale on the right). (D) Plot of the distribution of mean speed for NGN2 neurons at 3 days of terminal differentiation. (E) Plot of the distribution of distance traveled for NGN2 neurons at 3 days of terminal differentiation. (F) Plot of the distribution of directionality index for NGN2 neurons at 3 days of terminal differentiation. For D-F) Tracks that lasted less than 30min were excluded from the analysis (see Methods). *n*=5805 control NGN2 neurons, from 3 independent experiments. The mean values (dotted lines) are detailed on the plot for each condition ± SEM, while the darker areas show standard deviations. The plots were cut-off to better show the main populations, but the maximum values for mean speed and distance traveled are 2.9 µm/min and 390.8 µm respectively. (G) Plot of the distribution of nucleus aspect ratios for fixed NGN2 neurons at 3 days of differentiation. *n*=4095 NGN2 nuclei, from 3 independent experiments. The mean value (dotted line) is detailed on the plot ± SEM, while the darker area shows standard deviations. This plot was cut-off to better show the main population, the maximal value for nucleus aspect ratio is 5.5.

As with differentiation and neurite extension, our results on nucleokinesis compare favorably with other iPSC-based models and with rodent cultures (See Table 2), making NGN2 neurons an excellent model system for studying early stages of neuronal development and the underlying functions of the microtubule cytoskeleton.

**Table 2:**
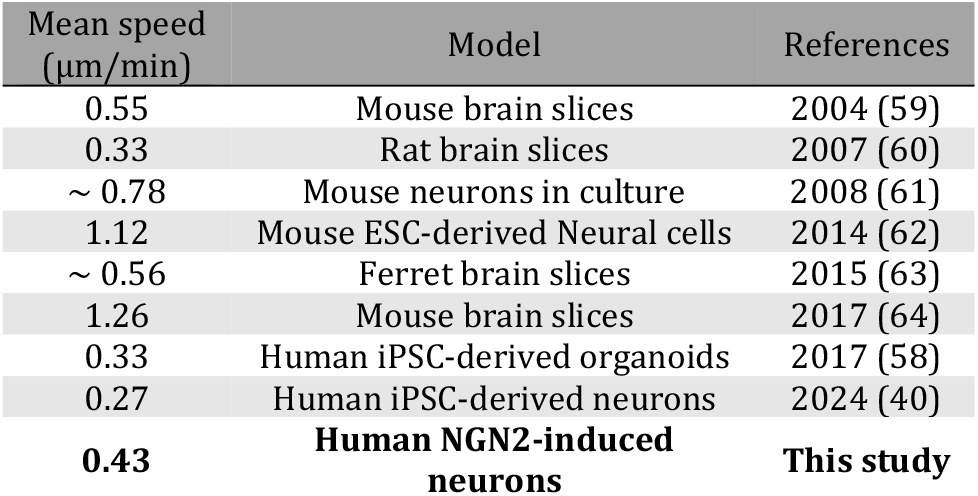
Nuclear mean speed in various models. Nucleus mean speed (µm/min) in different study models with references and date of publication. “∼” was used when values were extrapolated from multiple conditions or visually assessed from a graphical representation of the data.

| Mean speed ($\mu\text{m}/\text{min}$) | Model | References |
| --- | --- | --- |
| 0.55 | Mouse brain slices | 2004 (59) |
| 0.33 | Rat brain slices | 2007 (60) |
| ~ 0.78 | Mouse neurons in culture | 2008 (61) |
| 1.12 | Mouse ESC-derived Neural cells | 2014 (62) |
| ~ 0.56 | Ferret brain slices | 2015 (63) |
| 1.26 | Mouse brain slices | 2017 (64) |
| 0.33 | Human iPSC-derived organoids | 2017 (58) |
| 0.27 | Human iPSC-derived neurons | 2024 (40) |
| 0.43 | Human NGN2-induced neurons | This study |

### NGN2 morphology is perturbed by some, but not all, drugbased challenges

Both neurite extension and nucleokinesis require a sophisticated microtubule network divided into distinct, functional patterns. Thus, we wondered how NGN2 development would respond to perturbations to the microtubule cytoskeleton when challenged with four different drugs. Such perturbations serve to test the sensitivity and utility of our methods. Two drugs target microtubules directly (a taxane, paclitaxel; and a vinca alkaloid, the Diazonamide-A derivative DZ-2384), and two drugs target microtubules indirectly (an inhibitor of the severing enzyme spastin, spastazoline; and an inhibitor of a PTM pathway, trichostatin A). We incubated NGN2 neurons with each drug for 7 hours prior to analysis on day 3 in differentiation media. To analyze morphology and cell density, the neurons were fixed and stained for DAPI and Tubb3, and immunocytochemistry images were processed and analyzed as described above (Fig1).

Both drugs that target microtubules directly caused significant reductions in arborization lengths. Treatment with 100 nM paclitaxel caused the neurons to have shorter, curvier neurites, as if they were more flexible than control neurites (Fig. 3A). Paclitaxel binds to microtubules at their taxane sites, located in the lateral loops of β-tubulin in the microtubule lumen (*65*). This interaction creates structure in the lateral loops (*66*) and “expands” the microtubule lattice (*67*). These changes act to stabilize the microtubules but also to reduce their persistence lengths (*4, 9*). The 100 nM paclitaxel treatment did not appear to affect cell density, as measured by the number of nuclei per field of view (Fig. 3B), but treated neurons had a significantly reduced arborization length (Control: 203.7 ± 5.2 vs paclitaxel: 160.1 ± 6.0 µm, p<0.001, ***, Fig. 3C). We also observed a reduced intensity of Tubb3 staining (Fig. 3D), indicating either reduced expression of this isotype or overall reduced polymer levels (note: our fixation protocols include an extraction step that should remove soluble tubulin). The cells also covered a smaller area within the field of view (Fig. 3E), consistent with the reduced arborization length measured by skeletonization.

**Figure 3:**
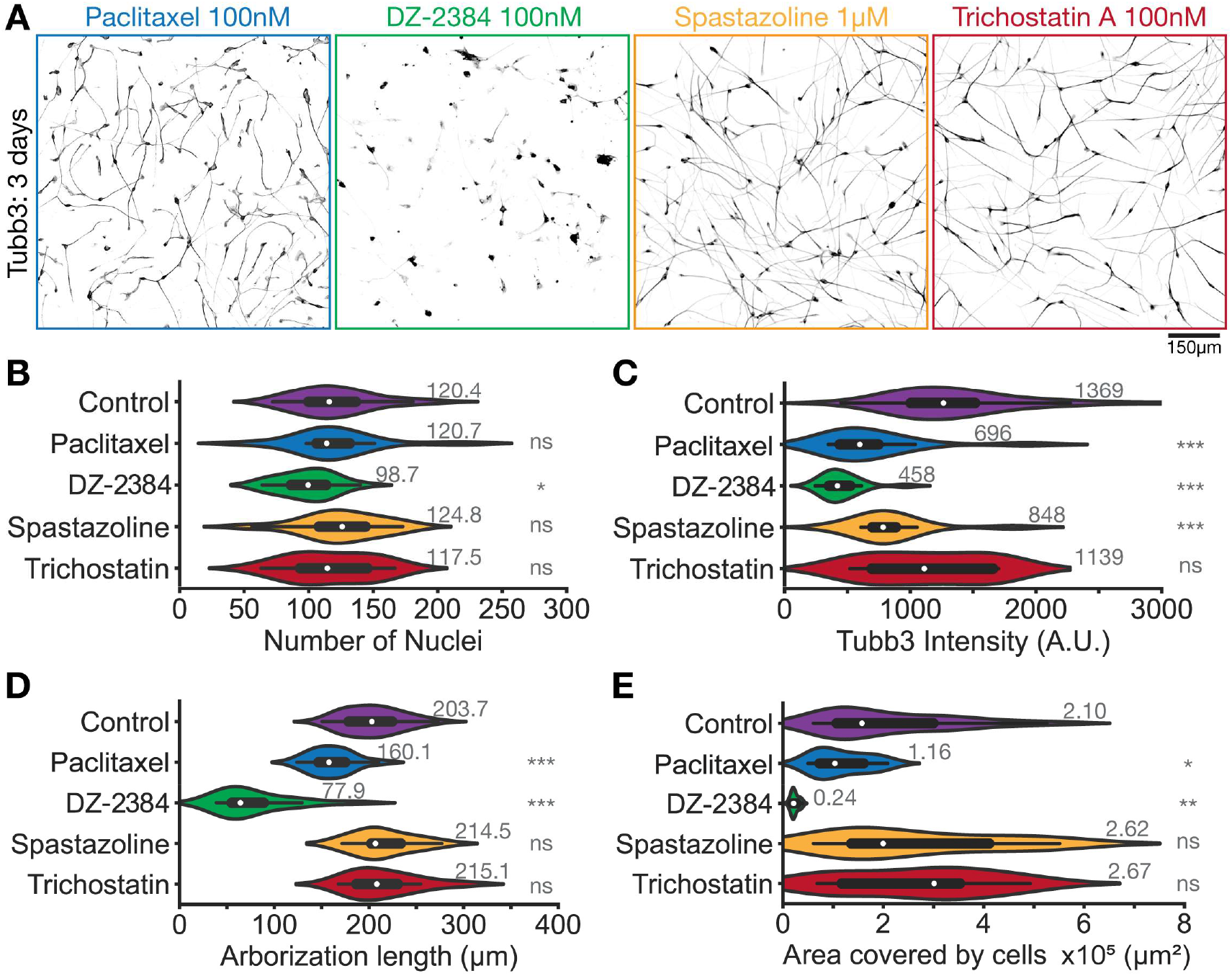
NGN2 morphology is perturbed by some, but not all, drug-based challenges. (A) Representative images of Tubb3 staining in 7h drug treated NGN2 neurons at 3 days of differentiation. The drugs were applied at the following concentrations: Paclitaxel 100nM, DZ-2384 100nM, Spastazoline 1µM, Trichostatin A 100nM. (B) Plot of the distribution of number of nuclei per field of view in control and drug treated NGN2 neurons at 3 days of differentiation. (C) Distribution of Tubb3 intensity per field of view in control and drug treated NGN2 neurons at 3 days of terminal differentiation. This plot was cut-off to better show the main population, the maximal value for Tubb3 intensity is 3156.4 A.U. (D) Plot of the distribution of neurite arborization length per cell body in control and drug treated NGN2 neurons at 3 days of terminal differentiation. *n*=1410-4095 neurons, from 3 independent experiments. (E) Plot of the distribution of the area covered by cells or Tubb3 staining in control and drug treated NGN2 neurons at 3 days of terminal differentiation. For B-E) Mean values are detailed next to the violins. On each violin, the white dot represents the median value, while the black bars show the quartiles/whiskers. *n*=12-34 field of views, from 3 independent experiments. Statistical analysis was done using Mann-Whitney non-parametric tests: ns non-significant, * *p*<0.05, ** *p*<0.01, *** *p*<0.001.

We observed similar, albeit stronger effects with DZ-2384 (*68-70*). Treatment with 100 nM DZ-2384 caused a sharp reduction in arborization length (Control: 203.7 ± 5.2 vs DZ-2384: 77.9 ± 9.8 µm, p<0.001, ***, see Fig. 3D), in Tubb3 signal (Fig. 3C), as well as cell area (Fig. 3E). DZ-2384 binds the vinca-alkaloid site, located at the interdimer interface of protofilaments, forcing the tubulin oligomers to adopt a curved conformation (*70*). Such oligomers prevent polymerization and/or promote destabilization of microtubule ends, typically leading to microtubule depolymerization, consistent with our findings. DZ-2384 also significantly reduced the number of nuclei per field of view (Fig. 3B), suggesting that this treatment causes cell detachment and/or cell death. These results demonstrate that direct disruptions to microtubule dynamics significantly changed neuronal morphology, regardless of the specific drug employed.

In contrast, spastazoline and trichostatin A, which target microtubules indirectly, did not affect neuronal morphology. We treated neurons with 1 µM spastazoline, an inhibitor of the microtubule severing enzyme, spastin (*71, 72*) and no morphological changes to neurons were apparent. Although its role(s) in neuronal development remain unclear, spastin localizes to neuronal branching points and has been proposed to sever long microtubules, thus generating short, dynamic microtubules that drive branch formation (*73, 74*). Alternatively, spastin may catalyze the turnover of tubulin within the microtubule shaft by creating damage in the lattice that is subsequently repaired (*75*). In either case, the 1 µM spastazoline treatment did not significantly change arborization length (Control: 203.7 ± 5.2 vs spastazoline: 214.5 ± 8.7 µm, *p*=0.33, ns, see Fig. 3D), cell density (Fig. 3B), or the area covered by Tubb3 signal (Fig. 3E). We did measure a reduced Tubb3 signal intensity (Fig. 3C), indicating the drug-based challenge was disruptive, but this reduced intensity did not translate into measurable morphological changes.

Finally, we challenged the microtubule network of NGN2 neurons by targeting a PTM pathway, namely acetylation. Acetylation occurs mostly at lysine 40 (*K40*) of α-tubulin, which is located in the microtubule lumen (*76*). The modification is reversible, and the levels of K40 acetylation reflect the balance of activity of acetylases (e.g., αTAT1) and de-acetylases. Once acetylated, microtubules are more stable and resistant to mechanical damage (*4, 9*). To increase acetylation levels, we treated NGN2 neurons with 100 nM trichostatin A, an inhibitor of histone deacetylase 6 (HDAC6). As its name indicates, HDAC6 can deacetylate histones as well as K40 α-tubulin (*76*). Similar to spastazoline, trichostatin A did not significantly affect arborization length (Control: 203.7 ± 5.2 vs trichostatin A: 215.1 ± 10.6 µm, *p*=0.45, ns, see Fig. 3D), cell density (Fig. 3B), or Tubb3 area (Fig. 3E), and no morphological changes were observed (Fig. 3A). We also did not observe changes in Tubb3 intensity (Fig. 3C). These results indicate that increased acetylation levels (confirmed by Western blots, Fig. S1) are not sufficient to modify the morphology of NGN2 neurons in a measurable way, at least under the conditions of our experiments. It remains possible that NGN2 neurons would respond to longer incubation times or an earlier challenge, but further experiments would be needed to completely exhaust these options.

Overall, these results confirm that disrupting microtubule dynamics directly with paclitaxel or DZ-2384 has a strong and rapid effect on NGN2 morphology. In contrast, increasing acetylation levels with trichostatin A or reducing Tubb3 signal intensity with spastazoline alone does not. Thus, the proper regulation of microtubule dynamics is critical in NGN2 neurons, as in many other model systems (*77*), and our methods are effective for demonstrating the impact of microtubule-targeting drugs on neurite extension.

### Changes in nuclear movements correlate with changes in morphology

Next, we wondered whether these changes in NGN2 morphology would correlate with changes in nucleokinesis. To that end, we analyzed NGN2 nuclear movements over a 5h time-period in the presence of our panel of drugs. Figure 4A and Figure 4B show representative images and tracking results for each condition, while Figure 4C shows a sample of extracted tracks (n=1322-1742 tracks per condition). As described previously, we measured the characteristics of these trajectories (e.g. mean speed); the distributions of these features are shown as violin plots in Figure 4D-G. Overall, each drug condition that we tested had a significant effect on at least some, if not all, characteristics of nucleokinesis.

**Figure 4:**
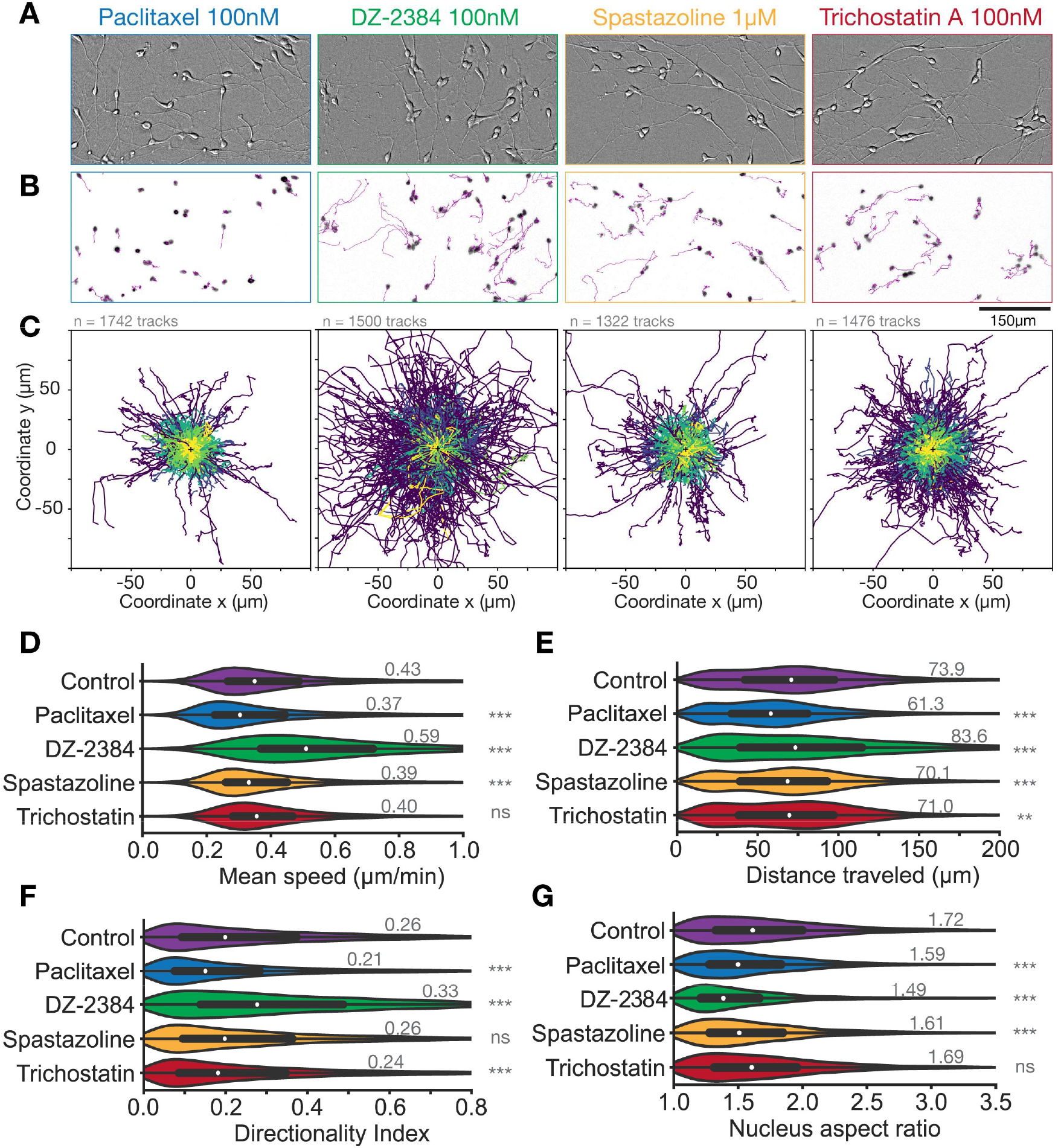
Changes in nuclear movements correlate with changes in morphology. (A) Representative still brightfield images of drug treated NGN2 neurons at 3 days of differentiation, showing neuronal morphology at the start of the tracking experiment. The drugs were applied at the following concentrations: Paclitaxel 100nM, DZ-2384 100nM, Spastazoline 1µM, Trichostatin A 100nM. (B) Representative still images of nuclei stained with Syto™ deep red for drug treated NGN2 neurons at 3 days of differentiation. Results of the automated tracking over a 5h-time period are showed as overlayed tracks for all conditions. (C) Position plots of 1742, 1500, 1322, and 1476 sample tracks for paclitaxel, DZ-2384, spastazoline and trichostatin treated NGN2 nuclei respectively. Each track is color-coded according to its displacement (see color scale Fig2C). (D) Plot of the distribution of mean speed for control and treated NGN2 nuclei at 3 days of terminal differentiation. (E) Plot of the distribution of distance traveled for control and treated NGN2 nuclei at 3 days of differentiation. (F) Plot of the distribution of directionality index for control and treated NGN2 nuclei at 3 days of terminal differentiation. For D-F) *n*=3859-5805 nuclear tracks from 3 independent experiments. These plots were cut-off to better show the main populations, maximal values are: mean speed 2.9 µm/min (Control), distance traveled 434.8 µm (Trichostatin). Statistical analysis was done using Kruskal-Wallis non-parametric tests: ns non-significant, ** *p*<0.01, *** *p*<0.001. (G) Distribution of nuclei aspect ratios for control and treated NGN2 neurons at 3 days of terminal differentiation. *n*=1410-4095 nuclei, from 3 independent experiments. This plot was cut-off to better show the main population, the maximal value for nucleus aspect ratio is 5.5 in the control condition. Statistical analysis was done using Mann-Whitney non-parametric tests: ns non-significant, *** p<0.001.

Similar to the analysis of NGN2 morphology (Fig. 3), the most visible differences in nuclear translocation were observed in the paclitaxel and DZ-2384 conditions. However, unlike the morphology results, these two direct microtubule-targeting drugs differed significantly in terms of their impact. Paclitaxel treatment inhibited nuclear movements, while DZ-2384 treatment seemed to exaggerate them. More precisely, paclitaxel treatment resulted in reduced mean speed, (Control: 0.43 ± 0.01 µm/min vs Paclitaxel: 0.37 ± 0.01 µm/min, p<0.001, ***, Fig. 4D), reduced distance traveled (Fig. 4E), and a lower directionality index (Fig. 4F). In addition, the nuclei were rounder than in control conditions (Fig. 4G), consistent with nuclei that are experiencing reduced cytoskeletal forces (reduced persistence length).

In contrast, DZ-2384 treatment resulted in higher mean speed (Control: 0.43 ± 0.01 µm/min vs DZ-2384: 0.59 ± 0.01 µm/min, p<0.001, ***, Fig. 4D), longer distance traveled (Fig. 4E), and increased directionality (Fig. 4F). Surprisingly, when looking at the nuclear shape, we found that DZ-2384-treated nuclei were rounder than control ones (Fig. 4G), suggesting there are no cytoskeletal forces applied to them. One hypothesis could be that microtubule depolymerization induced by DZ-2384 leads to detachment of neurite tips and subsequent retraction of neurites. These results emphasize the importance of both dynamic (enriched with DZ-2384) and stable (enriched with paclitaxel) microtubule populations for nuclear translocation and therefore neuronal migration.

Interestingly, whereas spastazoline had no effect on arborization length (Fig. 3D), its influence on nuclear movement was more pronounced. For example, spastazoline treatment resulted in reduced mean speed, (Control: 0.43 ± 0.01 µm/min vs spastazoline: 0.39 ± 0.01 µm/min, p<0.001, ***, Fig. 4D), reduced distance traveled (Fig. 4E), and a rounding of nuclear shape (Fig. 4G). This result is reminiscent of the data showing that disruption of centrosomal microtubules significantly affected nuclear migration in rodent cortical projection neurons (27). We can speculate that spastin may have a different impact on centrosomal vs. acentrosomal subsets of microtubules, although this idea requires testing.

Unlike the other drugs, trichostatin A had a minimal impact on both cell morphology and nucleokinesis. Trichostatin A treatment resulted in a small reduction in mean speed (Control: 0.43 ± 0.01 µm/min vs trichostatin A:0.40 ± 0.01 µm/min, *p*=0.16, ns, Fig. 4D), distance traveled (Fig. 4E) and directionality (Fig. 4F), but overall trichostatin A nuclei were the most statistically similar to control. These minimal effects on NGN2 nuclei support a hypothesis wherein acetylation of α-tubulin K40 is not relevant for nucleokinesis. Alternatively, we cannot exclude that phenotypes would manifest with longer drug treatments, although we do note that a significant increase in K40 acetylation is detectable in our time window (Fig. S1). Current models for the role of K40 acetylation suggest it plays a role in the regulation of intracellular trafficking, particularly pre-synaptic cargos (78, 79). However, such regulation may only become critical at later stages of development than those analysed here.

Altogether, our data indicates that NGN2 nucleokinesis is sensitive to a wider range of drugs than NGN2 morphology. The drug-based challenges to the NGN2 cytoskeleton suggest that the maintenance of microtubule patterns is critical to proper development in this model system. Indeed, having more stable (paclitaxel) or more dynamic (DZ-2384) microtubules both resulted in reduced arborization, and affected nucleokinesis although in opposite ways. Moreover, the spastin disruption by spastazoline implies some distinction between nucleokinesis and process extension, supporting the existing model that distinct subsets of microtubules are required for each of these developmental steps (*27*).

## Conclusion

We have developed open-access, semi-automated methods to measure the morphology and nucleokinesis of the latest generation of iPSC-derived neurons using doxycy-cline-induced expression of Neurogenin-2, which bypass the time-consuming process of inducing iPSCs into neural progenitor cells using SMADi. Terminal differentiation of NGN2 neurons into cortical neurons produced cultures that were homogenous and well-behaved; these neurons are perfectly tailored for drug screening studies and high-throughput data acquisition. Our study contributes to the build-out of this important model system through the comprehensive measurement of their process extension and nucleokinesis. We find that NGN2 neurons compare favorably to rodent primary cultures as well as previously described iPSC systems, and we are optimistic about their potential as a robust model system for the neuronal microtubule cytoskeleton. When the NGN2 cytoskeleton was challenged by drug compounds, the results underlined the importance of distinct patterns of microtubules in neuronal development (dynamic vs. stable, centrosomal vs. acentrosomal). We anticipate that iPSC-based cellular models will play a critical role in mapping out the molecular mechanisms by which neurons establish their complex cytoskeletons.

## Materials and Methods

### NGN2 iPSC-derived neurons culturing conditions

EBiSC-NEUR1 human-derived neurons (called NGN2 neurons throughout the paper) were obtained from the European Bank for induced pluripotent Stem Cells (EBiSC) repository (https://ebisc.org/EBISC-NEUR1). According to the provider’s specification sheets, the iPSC donor line BIONi010-C-13 (also available to purchase: https://ebisc.org/BI-ONi010-C-13, Sample ID: SAMEA103988285) comes from an African American male donor with no disease phenotype. At the repository location, the iPSC line was initially amplified as embryoid bodies in neurobasal media for 2 days then treated with doxycycline (#D3447, Sigma, 2µg/ml final) for 2 days, resulting in early neurons. These cells were dissociated and cryopreserved in Cryostor10 (#07930, Stemcell Technologies) in liquid nitrogen, then shipped to our lab. Once received, the cells were thawed rapidly in a 37°C water bath and seeded on coverslips (for immunofluorescence), dishes (for immunoblot), or glass bottom 96-well plates (for live imaging) that had been coated with PLO (#P3655, Sigma, 10µg/mL) and laminin (#L2020, Sigma, 5µg/mL). The cells were thawed and then grown for 3-7 days in “complete BrainPhys”: BrainPhys basal media (#05790, STEMCELL Technologies) supplemented with SM1 (#05711, STEMCELL Technologies, 2% final), N2A (#07152, STEMCELL Technologies, 1% final), BDNF (#78005, STEMCELL Technologies, 20 ng/mL final), GDNF (#78058, STEMCELL Technologies, 20 ng/mL final), cAMP (#100-0244, STEMCELL Technologies, 0.5mM final) and ascorbic acid (#A5690, Sigma, 200nM final). For the first 24 hours post-thaw, the cells were cultured in complete BrainPhys with 10µM Rock inhibitor. Doxycycline induction was maintained for 4 days post-thaw to ensure complete trans-differentiation. The media was changed daily until drug treatment, imaging, or fixation.

### Sample preparation, immunofluorescence and immunoblot

For all immunocytochemistry experiments, we used cells on their 3^rd^ or 7^th^ day of differentiation in complete BrainPhys. Treatments with micro-tubule-targeting drugs for staining were performed on the 3^rd^ day of differentiation for 7 hours. All drugs were diluted directly in complete BrainPhys right before treatment. Paclitaxel (#Y0000698, European Pharmacopeia Reference Standard), trichostatin A (#T1952, Sigma-Aldrich), and DZ-2384 (Paraza Pharma, (69)) were used at 100nM, while Spastazoline (#SML2659, Sigma-Aldrich) was used at 1µM. These concentrations correspond to 10-20x IC50 for each compound.

For immunofluorescence staining, cells were incubated in extraction buffer (PIPES 80mM, EGTA 1mM, MgCl2 7mM NaCl 150mM, D-glucose 5mM, Triton X-100 0.3%, glutaraldehyde 0.25%, (80)) for 1min30s at 37°C, before fixation in 4% PFA at 37°C for 15 minutes. The fixed coverslips were rinsed and preserved in PBS until staining. For staining, the samples were first quenched in NaBH4 (1mg/mL) for 10 minutes, rinsed in PBS, and permeabilized in PBS-Triton 0.5% for 10 minutes. Non-specific binding was blocked by incubation in blocking buffer (PBS-Triton 0.1%-BSA 3%) for 45 minutes. The coverslips were then bathed for 16 hours at 4°C in a mix of primary antibodies (listed below) diluted in blocking buffer. The samples were then rinsed in PBS-Triton 0.1%, before incubation for 2 hours at room temperature in a mix of fluorophore-conjugated secondary antibodies diluted in blocking buffer. After rinsing in PBS, DAPI stain was applied for 5 minutes at room temperature before mounting the coverslips on glass slides with Fluorsave (#345789, Millipore).

For immunoblots, media was removed from cell dishes and 2X Laemmli buffer (125mM Tris-HCL pH 6.8, glycerol 20%, bromophenol blue 0.04%, SDS 4%) freshly supplemented with DTT (100mM) was added directly to the plates. Cell lysates were scraped down and collected. These crude lysates were sonicated, centrifuged, and boiled for 5 minutes at 90°C before loading onto 4-12% acrylamide precast gels (#M41215, Genscript). Electrophoresis was performed in Tris-MOPS-SDS buffer (#M00138, Genscript) at 160V. Proteins were transferred to nitrocellulose membranes at 350mA for 90 minutes. Non-specific binding was blocked in TBST-Milk 5% (blocking buffer) for 45 minutes. The membranes were then incubated for 16 hours at 4°C in a mix of primary antibodies (listed below) diluted in blocking buffer. After 3 rinses in TBST, the membranes were incubated for 2 hours at room temperature with HRP-conjugated secondary antibodies diluted in blocking buffer. The membranes were rinsed in TBST again, and proteins were revealed using SuperSignal Pico PLUS ECL (#34580, Thermo) and exposed on film (#DIAFILM57, Diamed). Since native and modified tubulin have similar molecular weights, all films are from separate membranes of the same samples.

Primary antibodies were used as follows: anti-β3-tubulin (Tubb3, #802001, Biolegend) IF 1/500 -WB 1/10000; anti-acetylated α-tubulin (611B1, #sc23950, Santacruz Biotechnologies) IF 1/500 – WB 1/2000; anti-tyrosinated tubulin (YL1/2, #sc53029, Santacruz Biotechnologies) IF 1/50 – WB 1/1000; and anti-polyglutamylated tubulin (polyE, #AG-25B-0030-C050, Adipogen) IF 1/500 – WB 1/2000. Fluorophore-conjugated secondary antibodies were used at 1/500 while HRP-conjugated ones were used at 1/5000.

### Data collection

All microscopes used in this manuscript are part of the McGill University Advanced BioImaging Facility (ABIF), RRID:SCR_017697. The nucleokinesis dataset was acquired on a Zeiss Axio Observer Inverted Widefield Microscope, using the 20X Plan-Apochromat, NA=0.8 air objective. Cells were kept at 37°C/5% CO2 during imaging using a Live Cell Instruments CU-501 environmental control system. Prior to live imaging of NGN2 neurons on their 3^rd^ day of differentiation, Syto™ Deep Red nucleic acid stain (#S34900, Invitrogen, 250nM final) diluted in complete BrainPhys was applied to the cells for 20 minutes. The dye was then removed and replaced by complete BrainPhys at 37°C containing the microtubule-targeting drugs or vehicle. Images were taken every 5 minutes for 5 hours. At least 4 independent fields of view were recorded per well and each condition was duplicated in the same experiment. Immunofluorescence images were acquired on a Zeiss LSM710 inverted confocal laser scanning microscope using the 10X Plan-Neofluar NA=0.3 air objective. All images acquired in NGN2 neurons with or without drugs were acquired in parallel with the same laser power and gain conditions, to allow comparison of the independent experiments.

### Analysis

Fiji was used for all image processing and data measurements and Microsoft Excel and Python were used for graphical representation of the data. All macros and scripts used in this study are available on GitHub (https://github.com/brouhardlab/Sebastien-2024).

#### Nuclei tracking

An automated TrackMate-based pipeline (*56, 57*) built in Fiji was used to generate nuclei tracks for each field of view. For identification of the nuclei througout timelapses, the Laplacian of Gaussian (LoG) detector was used since it best detected Gaussian-like structures. A 10 µm diameter setup was used based on initial measurements of nuclear size. For linking the detected spots between frames, the simple Linear Assignment Problem (LAP) tracker was used as the simplest and most effective algorithm. A subpixel detection was applied to avoid jagged trajectories and the other linking parameters were defined empirically on a few test images. We used a 20 µm maximum linking distance and gaps of 4 frames maximum were allowed, and the background value was assumed to be below 5.0 A.U. Tracks and spots data were exported as .xml files, while processed trajectories were exported as .csv files. The track .xml files were used for visual representation of trajectories (Fig. 2 and 4). A range of information can also be extracted from the tracks and spots data, e.g. intensity of the different channels and precise (*x,y*) coordinates. The mean speed, distance traveled, displacement, and directionality index (Displacement divided by distance traveled (*40*)) were obtained from the .csv file. After visual assessement of the accuracy of the tracking data on each file, all processed .csv files were compiled in Python for graphical representation (Python script “Nuclear tracking”). Only tracks that lasted more than 30 minutes were included in the quantifications (75% of all tracks), because shorter tracks were most often found to be tracking errors and false-positives.

#### Neurite extension

To measure the average arborization length per cell, we used the Tubb3 and DAPI immunofluorescence stainings in a homemade Fiji macro (Macro “Neurite arborization”) and we iterated through the dataset via Fiji’s “batch” function. To determine the total branch arbor in an image, we first removed signal heterogeneity by applying a pseudo-flatfield correction to the Tubb3 channel. The channel was subsequently passed through a gaussian blur filter (sigma=1µm) and binarized with Otsu thresholding. Then the image was skeletonized and the fraction of the image (“Area”) covered by the skeletonized neurites was measured. Since the skeletonization resulted in lines with a single pixel width, the area value was used as a total length value. In parallel, the DAPI channel was filtered using a Gaussian blur (sigma=1 µm), then binarized with Otsu thresholding. The number of nuclei in the image was detected with the “Analyze particles” function with a particle size of 25-150 µm^2^ and circularity of 0-1. To obtain a value of average neurite extension per neuron, we divided the total neurite length (“Area”) by the number of nuclei in the image.

#### Tubulin intensity and area

Since all conditions were seeded together, the cell density in each image is considered the same for all treatments, we therefore used Fiji macros again with the batch function to measure the tubulin intensity in whole field of views (Macro “Intensity FOV”), and then the area covered by the cells in these images (Macro “Int-Area”). First, the Tubb3 intensity was measured in the full image. To get the area covered by the cells, the Tubb3 channel was passed through a pseudo flat-field correction before binarizing with Otsu thresolding. The “Analyse particles” function was then used with particle size of 50-Infinity (µm^2^) and circularity of 0. The obtained ROIs were combined as one and this area were measured. The ROIs can be further applied on the raw image to measure the signal of any channel inside the cells (Signal density).

#### Nucleus AR

The aspect ratio for all nuclei was measured from the DAPI channel. As previously, the DAPI channel was filtered with a gaussian blur (sigma=1) and binarized using Otsu thresholding. We then used the “Analyze particles” function and the “Shape descriptors” measurement to obtain an aspect ratio for each nucleus.

### Statistics

All statistical tests used are indicated in the figure legends: Kruskal-Wallis non parametric H test, or Mann-Whitney non parametric U test. P values were considered nonsignificant when above 0.05. In some plots, the axes were truncated for display purposes; the figure captions clearly indicate such cases.

## Acknowledgements

We would like to thank Isabelle Sébastien, Jeanette Wihan and Julia Neubauer from the Fraunhofer IBMT, for their excellent support and guidance on the use of NEUR1 neurons throughout the project. We would also like to acknowledge the ABIF facility for their helpful support with data collection. GJB acknowledges support from Natural Sciences and Engineering Research Council of Canada (RGPIN-2014-03791), the Canadian Institutes of Health Research (PJT-148702), and McGill University. AGH is supported by the Canadian Institutes of Health Research (PJT-185997).

## Author contributions

Conceptualization and experimental design: MS and GJB; Experimental procedures and data analysis: MS and AP; Code writing: AGH, LF and MS; Manuscript writing: MS and GJB; Manuscript editing: all authors; Funding acquisition: GJB and AGH.

## Supplemental figures

**Figure S1:**
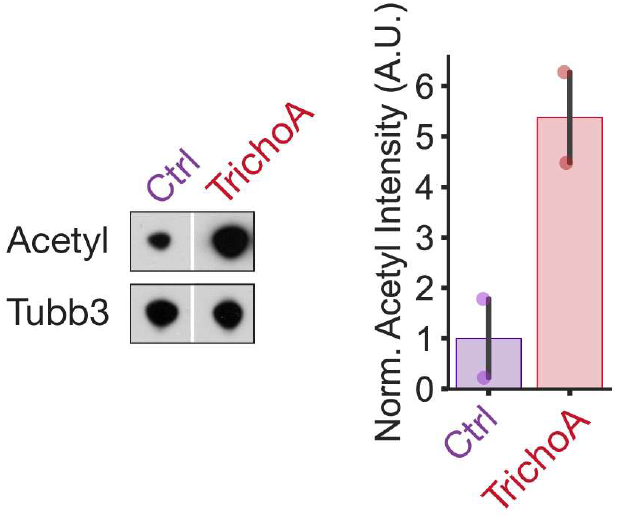
Trichostatin A treatment increases acetylation level. Representative immunoblot and quantification showing the increase in acetylation upon treatment with Trichostatin A. Control and Trichostatin samples are on the same membrane, and they were exposed together for the same amount of time.

